# Improving modelling for epidemic responses: reflections from members of the UK infectious disease modelling community on their experiences during the COVID-19 pandemic

**DOI:** 10.1101/2023.06.12.544667

**Authors:** Katharine Sherratt, Anna C. Carnegie, Adam Kucharski, Anne Cori, Carl A. B. Pearson, Christopher I. Jarvis, Christopher Overton, Dale Weston, Edward M. Hill, Edward Knock, Elizabeth Fearon, Emily Nightingale, Joel Hellewell, W. John Edmunds, Julián Villabona Arenas, Kiesha Prem, Li Pi, Marc Baguelin, Michelle Kendall, Neil Ferguson, Nicholas Davies, Rosalind M. Eggo, Sabine van Elsland, Timothy Russell, Sebastian Funk, Yang Liu, Sam Abbott

## Abstract

The COVID-19 pandemic both relied and placed significant burdens on the experts involved from research and public health sectors. The sustained high pressure of a pandemic on responders, such as healthcare workers, can lead to lasting psychological impacts including acute stress disorder, post-traumatic stress disorder, burnout, and moral injury, which can impact individual wellbeing and productivity. As members of the infectious disease modelling community, we convened a reflective workshop to understand the professional and personal impacts of response work on our community and to propose recommendations for future epidemic responses. The attendees represented a range of career stages, institutions, and disciplines. This piece was collectively produced by those present at the session based on our collective experiences. Key issues we identified at the workshop were lack of institutional support, insecure contracts, unequal credit and recognition, and mental health impacts. Our recommendations include rewarding impactful work, fostering academia-public health collaboration, decreasing dependence on key individuals by developing teams, increasing transparency in decision-making, and implementing sustainable work practices. Despite limitations in representation, this workshop provided valuable insights into the UK COVID-19 modelling experience and guidance for future public health crises. Recognising and addressing the issues highlighted is crucial, in our view, for ensuring the effectiveness of epidemic response work in the future.

## Introduction

The response to the COVID-19 pandemic necessitated a multi-pronged approach, with infectious disease transmission modelling playing a key role in informing strategy and policy decisions [1, 2]. Input from UK modellers was mostly channelled through weekly meetings of the Scientific Pandemic Influenza Group on Modelling, Operational subgroup (SPI-M-O) feeding into the Scientific Advisory Group for Emergencies (SAGE) [3]. This advisory group, drawing on expertise from the academic, and public health sectors, developed planning scenarios and short-to-medium term forecasts and projections, routinely estimated key parameters such as the reproduction number (a proxy for transmissibility), conducted routine data analysis, as well as authoring ad-hoc reports on modelling results relevant to the ongoing pandemic in the UK [4], [5]. Some of these analyses resulted in academic papers along with those produced by the wider UK modelling community (e.g. [6, 7]).

The high-pressure environment and daunting responsibilities of those at the frontlines of pandemic response have been shown to exert significant psychological tolls. Notably, healthcare workers (HCWs) involved in infectious disease outbreaks, including COVID-19, have been shown to experience profound and enduring psychological impacts. These include acute stress disorder, post-traumatic stress disorder (PTSD), burnout, as well as moral injury [8–10]. Moral injury refers to a specific form of distress that stems from guilt, anxiety, and loss of trust when actions or roles conflict with one’s deeply held moral beliefs. These psychological impacts not only diminish individual wellbeing but can also considerably affect worker productivity, with lasting effects that can linger for years, as exemplified by the 2002/2003 SARS epidemic [11], [12].

However, the experiences and challenges faced by non-healthcare responders to the pandemic, such as those involved in modelling and research, have received comparatively less attention [8, 10]. Stressors such as high workloads, long hours, tight deadlines, and harassment from the public and press during the COVID-19 response had the potential to cause both visible and invisible impacts. These include mental health impacts, exhaustion, social isolation, compromised career progression in academia, and moral injury.

The experiences of modelling responders have not been systematically discussed but are indirectly reflected in issues of staff retention and burnout across institutions. With the aim of bridging this gap, on March 28th 2023, we organised a one-day workshop to create a space for collective reflection and strategising improvements for future epidemic responses. This paper seeks to provide an outline of the workshop proceedings, the collective themes that emerged from our discussions, and synthesise our suggested actions into a set of priority recommendations to enhance future epidemic responses.

## Our approach

We employed an iterative, participatory approach to both design and run a reflective workshop with members of the UK modelling community in order to facilitate the summarisation of our collective experiences. In the interest of clearly relaying the proceedings and results emanating from the workshop, we use the term ‘participants’ to refer to attendees (i.e. ourselves, including the organisers) in the remainder of the methods and the results.

### Workshop design

We aimed to ensure the content of the reflective session captured the needs of the individuals at the forefront of the UK modelling response. To inform the content of the workshop, an online survey was developed by the session organisers (SA and ACC) alongside two additional members of the UK modelling community. The survey allowed free-text responses to questions about the personal and professional ramifications of participating in COVID-19 response work, along with the obstacles to effective response work and strategies to address them (see the Supplementary Information). The survey was circulated among members of the UK modelling community, advertising through organisation mailing lists and social media. Responses were anonymous. The survey was open between mid-December 2022 and mid-January 2023 and received 27 responses.

We then engaged an external facilitator to assist in planning the agenda and guiding participants throughout the session. This aimed to ensure unbiased management of discussions and to enable participants to express themselves openly in a safe and supportive environment. To select an appropriate facilitator, we sought input from the broader scientific community and chose an individual with a track record of successfully delivering similar events.

Initial discussion topics were developed by the session organisers in consultation with the external facilitator, drawing on the responses received from the survey. Further feedback was solicited from two members (KS and YL) of the UK modelling community who were not directly involved in the organisation process. This resulted in a set of discussion topics that addressed the concerns and interests of the survey respondents (see supplementary information).

### Participants

We aimed to include a diverse range of participants involved in the UK COVID-19 modelling response, encompassing researchers and professional services staff. A brief expression of interest form was disseminated by the session organisers to the UK modelling community via organisation mailing lists, personal networks (aiming to also reach those who may have transitioned away from the infectious disease modelling field), and social media channels to ensure representation across different levels of seniority. We invited all those who expressed an interest to attend. We provided a small travel fund for participants on a first come first served basis for those travelling from across the UK.

### Workshop structure

#### Participant arrival and introduction

Upon arrival, the facilitator encouraged participants to engage with flipchart papers displaying “snapshot” questions with attendees providing their responses using stickers. These were:

1. Do you think sufficient action is currently being taken to improve future outbreak responses to the standard you think is acceptable?
2. Who is responsible for ensuring people are supported, and appropriately credited for their work?
3. Summarise your pandemic experience in one word.

See the supplementary information for the multi-choice answers.

At the formal start time, the facilitator opened with an overview of the day’s agenda, establishing expectations and a code of conduct for participants. The Chatham House Rule (“share the information you receive, but do not reveal the identity of who said it” [13]) were introduced to ensure that individuals would not be identified, while allowing for the synthesis of outputs. A co-organiser (SA) shared their personal pandemic timeline (see the supplementary information), setting the stage for the first exercise.

#### Iterative discussions of experience

Participants divided themselves into pairs, after being encouraged to work with someone they would not usually interact with. They were asked to discuss their individual pandemic timelines for 15 minutes each, while the partner asked questions based on those we developed when designing the workshop, listened, and asked follow up questions. The following questions were provided.

1. What was your pandemic timeline? What were the highs and lows?
2. What was your experience of pandemic work like?
3. What were some of the things that helped assist you to do effective research during the outbreak response?
4. Do you think team science was appropriately supported over the pandemic?
5. Has your employer or the wider community taken action to help mitigate any of the personal or professional costs/challenges you identified? What more can be done?
6. Do you think there were barriers to doing effective and sustainable COVID-19 outbreak response work? If so, what were they?
7. What has been done and what more can be done to reduce any barriers to effective outbreak response work in the future?

We also provided suggested follow-on questions which are available in the supplementary information.

Pairs then formed groups of four to identify common themes from their one-to-one reflections using post-it notes. The group was then brought together and themes were summarised and organised into headline categories. This approach maintained anonymity for the participants, while capturing their reflections in a summarised form. As a combined group we then further discussed these topics, leading to the identification of six major themes.

#### Synthesising recommendations

The latter portion of the workshop focussed on pinpointing recommendations for action. Participants were presented with primary categories derived from the morning’s discussions. Participants were then divided into new groups, with each group assigned a theme. Each group was tasked with developing recommendations and potential implementers. Participants could move between themes and contribute their thoughts. These recommendations, along with actionable steps and suggestions for those responsible for implementing, were then exhibited on a wall for group review. Finally, attendees used dot stickers to identify priorities, allowing a visual representation of the group’s consensus.

We (ACC, KS, YL, SA) then reviewed the contents of the group discussions based on the post-it notes, whiteboards, and recommendation board created during the session. Two authors (ACC and KS) independently digitised the output, and four authors (ACC, KS, YL, SA) independently reviewed results. We then came to a consensus on the common themes of participants’ experiences, using the major themes identified by participants as a guide, and priority recommendations for stakeholders. We prepared an initial draft and shared this with participants. Finally, we integrated feedback, ensuring that the insights derived from the workshop were preserved.

## Outputs from the workshop

### Summary of attendees

The event was attended by 27 individuals, including 25 research staff and two professional services staff. Staff attended from five higher education institutions (London School of Hygiene & Tropical Medicine, Imperial College London, University of Warwick, Liverpool University, the University of Oxford), and the UK Health Security Agency (UKHSA). The majority of attendees were based in London. Participants represented various career stages, including early, mid-career, and senior academics and professionals. Among the attendees were multiple members of SPI-M-O and SAGE.

### Snapshot reflections

In response to the initial snapshot question, “Do you think sufficient action is currently being taken to improve response work to a standard you think is acceptable?”, the participants expressed an overwhelmingly negative view (17/18). In the second snapshot question, “Who should ensure that individuals are adequately supported and credited for their response work” (this question allowed multiple answers), participants suggested this responsibility was shared among stakeholders. Affirmative responses were more common on the panel listing smaller groups than on the panel listing larger organisations, indicating that respondents considered themselves (9/56), line managers (16/56), and research groups (12/56) more responsible for this task compared to larger organisations such as institutions (6/56), academic funders (5/56), and the “system more generally” (8/56). The second panel displaying the larger organisations was situated to the right of the first panel, which may have resulted in decreased visibility. Photos of the panels are available in the supplementary information.

Each attendee was asked to summarise their pandemic experience in one word using post-it notes (see supplementary information). Positive responses were: “*exciting”, “valuable”*, and “*engrossing”*. Neutral responses were: *“intense”, “unprecedented”, “ambiguous”, “focussed”, “hectic”,* “*repetitive”,* and “*surreal”*. Negative responses were: “*hard”, “austere”, “stressful”, “lonely”, “harrowing”, “frustration”*, and “*exhausting”*. Multiple participants added stickers (indicating support) to both “*stressful”* and “*exhausting”*.

### Themes from paired and group discussions

In the paired and small group discussions, several topics emerged into which participants’ perspectives were grouped. These included (reproduced verbatim from those listed on the day): *“societal impact”, “mental health”, “life outside”, “emotions”, “personal”, “team spirit”, “institutional structures”, “work process”, “work feeling/support”, “work pressure”, “(negativity about)” “positives”*, *“career direction”*, *“rewards”*, *“access and privilege”*, *“bad stuff”*, *“COVID-19 modeller-specific experiences”*, and *“general experiences”*. In the following session, participants then refined these themes to leave the following: *“institutional factors”, “mental health”, “life/personal”, “work process”, “career direction”, and “social impact”*. See the Supplementary Information for the full list of participants’ points.

After the workshop, we reviewed individual post-it notes and further refined these themes to leave: *funding and institutional support*; *recognition, rewards, and access*; *team and work dynamics*; *non-academic contributions*; and *personal impacts*. The themes emerging from the group discussions are synthesised, stratified by these themes below. We indicate direct quotes from individual authors using quotation marks and italics.

#### Funding and institutional support

*Lack of institutional support:* Insufficient institutional support for those involved in the COVID-19 modelling response was a common issue among participants. Many felt that they were not protected by their institutions during the response or in its aftermath, for example when receiving aggression from some sectors of the media and general public. Additionally, groups highlighted the lack of processes to respond in an emergency while protecting psychological safety. This included the need for training for managers and teams, and wellbeing procedures and Human Resources policies.

*Contract insecurity and inflexible funding rules:* The precarity of short-term contracts due to heavy reliance on external grant funding was highlighted, along with implicit pressures to underestimate personnel time in funding applications to meet budget thresholds, adhere to eligibility criteria and achieve cost recovery targets. The importance of providing sufficient and sustainable personnel funding was stressed, with this including academic and professional services roles such as project managers, administrators, communications professionals, technicians, and software engineers.

#### Recognition, rewards and access

*Inadequacy of reward metrics:* Credit attribution mechanisms were a recurring concern. Participants emphasised that there are currently insufficient frameworks to reward the nature of response work itself. Hurdles in receiving recognition for work included contributing to confidential reports where involvement was unable to receive external acknowledgement. In particular, it was noted that outputs such as software tools and policy reports do not fit within the traditional academic credit structure. Similarly, participants recognised that promotions, paper authorship, and grant Principal Investigator (PI) positions were not designed to promote collaborative team working. This was identified as a problem for both the general wellbeing of researchers and the quality of the science produced. The unequal, and individual-focussed, credit structures that persisted throughout the pandemic were also discussed, with senior or well-connected researchers being identified as receiving the majority of recognition. Participants noted *“rewards not attributed equally”,* and that *“institutions got awards, not individuals (not all key players)”.* This uneven reward system was seen as contributing to a competitive culture, which was identified as a problematic aspect of response work and academia more widely.

*Access to decision-making spaces:* Individuals had different access to policy-making spaces which did not always reflect where or how their work was used. As a result, some individuals who lacked access reported feeling left behind when it came to updates relevant to their work. There was a general consensus that there should be more transparency regarding these forums for those involved in producing the work presented.

#### Team and work dynamics

*Insufficient capacity:* Participants highlighted issues with *“not being able to say no”* and the *“pressure [that] came in waves. ‘Not again…’”* These issues contributed to poor working practices within teams, including insufficient capacity and reliance on one or two individuals to perform key tasks. In turn, this made it more challenging for these individuals to maintain a work-life balance. The highly pressurised and reactive nature of response work meant that there was not always space for teams to reflect on the effectiveness of routine aspects of the response, including whether academic groups were the best placed to perform this work. In addition, despite a need for additional capacity, working in highly reactive ‘response mode’ made it difficult to properly onboard new starters and hand over responsibility of tasks and projects where resources were available to do this. There was reference to other professions more adapted to response work, such as the military and emergency services, suggesting there may be learning to be gained from these sectors.

*Competing demands and barriers to progression:* Individuals faced challenges in balancing competing demands of and distinguishing between ‘response’ work and research. Some individuals sacrificed otherwise beneficial opportunities, such as teaching. Although response work created some opportunities for career progression, these were distributed unequally relative to contribution. Access to these opportunities depended on several factors, including career stage, and relative privilege (which is the differential access to resources, opportunities, and advantages some groups have compared to others). We note that privilege is often invisible to those who have it, and recognizing one’s own relative privilege is a key part of understanding and addressing social inequalities.

*Collaborative working:* Participants cited the positive experience of collaborative working and camaraderie within teams - academic, professional, and hybrid. However, as the pandemic progressed, there was a sense that the egalitarian working structures which some felt were put in place at the start of the pandemic faded: *“Shift from egalitarian structure to pre-existing hierarchies”*. Meanwhile, with close working relationships and the intensely personal impact of the COVID-19 response, professional disagreements sometimes took on an unusually emotional tone.

#### Non-academic contributions

*Role of professional services staff:* Participants highlighted the significance of integrating professional services roles into research teams, mentioning that these staff played crucial roles in response-related tasks. Participants pointed out that professional services roles, especially administrative positions like project managers, are frequently deemed ineligible costs in grant applications. Similar to academic staff, many individuals in these roles work on short-term contracts. Consequently, these positions were often under-resourced and experienced high turnover.

*Public health agency workers:* Participants emphasised the importance of strengthening the collaboration between academics and public health agencies, with the aim of fostering knowledge and skills exchange both during, prior to, and after responses. The importance of a bidirectional exchange was highlighted, with academics having the opportunity to learn about the practical challenges faced by public health agencies, while public health staff would benefit from access to the latest research findings. Participants called for more opportunities to facilitate these exchanges, such as joint workshops, shared working spaces, and dedicated training sessions.

#### Personal impacts

*Public recognition:* The COVID-19 pandemic brought the infectious disease modelling field public recognition and scrutiny. Participants acknowledged the personal responsibility that came with this visibility, while valuing the significance of their work. While friends and family gained deeper understanding of their work, some highlighted the challenge of work and life becoming intertwined. Participants referred to the *“surreal level of public and media interest (good or bad)”* and the idea that *“work and the world were one and the same. Neither was an escape from the other.”*

*Mental ill health and burnout:* Participants across organisations and seniority levels reported prioritising work over their health and wellbeing, leading to extreme levels of overwork, burnout, and associated mental health effects, including depression and anxiety. The experience was common among attendees at all levels and career types, with recognition that this can creep up over time and not enough has been done to mitigate against it. Some participants expressed guilt and a sense of ‘survivor bias’ from being able to remain within academia, having witnessed friends and colleagues leave the field. One post-it note summed up the feeling of *“trading off career versus health and everything”*. People were reluctant to reach out to managers or colleagues for support. With close working relationships, the personal challenges faced by colleagues inevitably impacted the wider team. No strategies were identified by participants as having been in place to address these issues during the pandemic response or having been implemented more recently.

*Commitments outside of work:* Several participants highlighted the challenges they faced in balancing high-intensity roles with personal obligations during the pandemic response. They shared experiences of coping with loss and caregiving responsibilities, which were particularly difficult for those whose partners were also involved in the response. Certain groups faced heightened challenges: for example, women often bore a disproportionate burden of caregiving tasks, early career researchers tended to have less stable domestic situations, and non-UK nationals experienced difficulties such as visa concerns or being separated from their home countries.

### Recommendations

The strategies collectively proposed at the workshop spanned societal impact, mental health, career direction, work processes, personal life, and institutional policy. Over ninety suggestions were made for possible actions by research teams, employers, and funding entities. The full list of recommendations is available in the supplementary information.

### Priority recommendations

Participants distilled a set of priority recommendations to enhance the support and sustainability of epidemic response work. These directives tackle crucial facets affecting the well-being and efficiency of those engaged in pandemic response. Example actions for implementing these recommendations are italicised below each recommendation (see the full list of suggested actions from the workshop in the supplementary information).

1. Acknowledge, and reward, impactful response work at institution, funder, and research community levels. *Funding bodies refine impact measures to credit all forms of output produced during, and required for, response work; institutions standardise incorporating response-driven work into criteria for doctoral theses and promotion*.
2. Encourage routine interaction between academia and public health agencies, including consistently reviewing the role of each during epidemic responses. *Government bodies and research institutes create sustainable dual positions recruiting from both sectors*.
3. Ensure response teams are well-staffed, well-resourced, stable, and provided psychological support. *Research teams establish sustainable team-building and training programmes with long term support from funders during non-response periods to ensure individuals feel equipped and supported to engage in response work*.
4. Increase the transparency of the evidence pathway from scientists to decision-makers making it easier for those across the scientific community to contribute as well as making the evidence base for decisions clearer to the general public. *Government bodies standardise rapid open access to the minutes of scientific advisory meetings and encourage input from a wider range of sources*.
5. Implement best practices for a sustainable work environment. *Employers promote leave-taking and respecting work hours, and clarify communication about processes and rewards across career stages, integrate support roles into research teams, and standardise the onboarding of new team members*.

## Discussion

This reflective workshop brought together 27 individuals from the UK infectious disease modelling community to engage in a dialogue around the personal and professional impacts of their COVID-19 response work. Participants represented various career stages, institutions, and disciplines, enabling a diverse exploration of experiences and perspectives. We identified areas of improvement in the current approach to modelling during epidemic responses, with these including greater support for responders, line managers, and research groups. Our experiences ranged from positive to negative, with stress and exhaustion being particularly prominent. Through in-depth discussions, key themes emerged, including institutional support, mental health, career direction, and social impact. Challenges such as lack of institutional backing, insecure contracts, inadequate reward systems, and personal impacts such as mental health issues were identified. The roles of professional services staff and public health agency workers were underscored. To address these issues, we identified a variety of strategies and priority recommendations, including acknowledgement and reward of impactful response work including for professional service staff, enhanced academia-public health collaboration, minimising dependence on key individuals, increased transparency in decision-making processes, and the adoption of sustainable work practices. These findings offer valuable insights for the ongoing pandemic response and future public health emergencies.

Our approach benefitted from being embedded in the experience of the UK modelling community. The session was community-driven, adopted an informal approach, and included participants from various career stages and perspectives on the response. Prior input from the community through an informal survey ensured the event’s relevance for attendees, while employing an external facilitator helped create a safe and structured environment for discussion. We then collectively agreed on key themes and recommendations.

However, a key limitation was participant representation. This was exacerbated by it being a one-day workshop, meaning we could only represent the views of those who were available and able to attend in person on that day. Attendees were primarily from London and South East England, possibly due to limited support for travel costs. Additionally, despite efforts to involve individuals who had left the infectious disease modelling field, few were able to attend. Our collective experiences are therefore likely to be missing some of the most challenging experiences and perspectives of responders, and our conclusions may be more moderate than if a wider range of participants had attended. Despite this bias, we feel this provides valuable insights into the UK COVID-19 modelling experience, but should be viewed as a summary of a small group’s experiences and opinions, with potential differences across jurisdictions and groups. We encourage responders in other locations to conduct similar exercises and to synthesise these findings for a broader understanding.

Ongoing efforts have begun to evaluate UK modelling work during COVID-19 both in terms of modelling results (e.g. forecasts or scenario projections [6, 7]), and the systems and processes enabling the response [2, 14, 15]. However, so far little has been done to report the experiences of responders themselves as we have done in this work. In the context of more general crisis response, more work has been done to understand the key challenges, particularly on healthcare workers (HCW). For example, hospital disaster preparedness plans may incorporate mental and behavioural health interventions (such as resource signalling, peer support, and referrals for at-risk individuals), which have proved to be effective in reducing mental health morbidities [16, 17]. Lessons from previous epidemics also emphasise the importance of effective staff support and training in preparing for future outbreaks. Perceived adequacy of training and support had a protective effect on adverse outcomes in HCW responders to the SARS epidemic [18]. These approaches, which have established use in high-stress occupations, could be adapted and applied to support modellers during epidemic response situations.

The workshop identified priority recommendations aimed at enhancing support and sustainability in pandemic response work. Our discussions underscored the importance of recognising and rewarding significant contributions to public health crises at all levels. We advocated for fostering closer ties between academia and public health agencies, building well-resourced, resilient teams, and ensuring their psychological well-being. Discussions also emphasised the need for increased transparency in the evidence-to-policy pipeline, improved work-life balance, and clear institutional communication. Further suggestions included standardising onboarding procedures and integrating support roles into teams.

Whilst we identified several themes and recommendations during our workshop, we did not explicitly separate issues specific to the pandemic response from broader academic challenges. Some recommendations, for example, recognising non-traditional contributions or normalising annual leave, pertain to broader issues. It is important to discern whether these concerns are long-standing systemic issues that have been simply exacerbated by the pandemic, or if they have been particularly highlighted due to the unique stressors of the pandemic.

## Conclusions

As a community we want to acknowledge that the pandemic has engendered widespread hardship, stress, and ill health throughout various populations. It is crucial to reflect on and address these profound impacts as we continue to tackle the crisis and prepare for future epidemic responses.

The consequences of the COVID-19 pandemic have been profound on those at the forefront of the UK modelling response. In this work, we have summarised our experiences and whilst we recognise that many of the issues we have identified impact those in our field more generally we believe that they are particularly problematic for epidemic response work. It is evident that changes are required across multiple domains, including individual work, team dynamics, and institutional structures, to enable future effective epidemic modelling responses.

Achieving these changes necessitates investment from governments, funding bodies and institutions. The solutions needed to foster a healthy and sustainable environment for future epidemic response work will not be attainable without such investment. Additionally, there is a need for teams aiming to respond to epidemics to redefine their working methods, developing response preparedness plans that emphasise wellbeing, training, and career development. It is clear that even these localised initiatives demand time investment from those leading them, and as a result, require support.

As it stands, future epidemic responses are likely to raise similar challenges to those we have identified here, including reliance on a select number of individuals, excessive workloads and the exacerbation of systemic inequalities. It is critical we act outside of response contexts, for example by implementing the recommendations we have outlined, to mitigate these issues and respond more effectively in future.

## Acknowledgements

We wish to acknowledge the support of the following individuals in the development of this project: Rosanna Barnard, Ciara Dangerfield, Dale Weston, Anne Cori, Joel Hellewell, Charlotte Hall, Rosalind Eggo, Stefan Flasche, Sebastian Funk, Mark Jit, Graham Medley, and external facilitator, Janice McNamara.

We also wish to thank everyone who so graciously shared their experiences at this event and those that completed the online survey.

## Supplementary Information

### Survey questions

How best would you describe your role in responding to COVID-19?

- Academic researcher
- Civil servant
- Professional services staff
- Other

How best would you describe the role you are doing now?

- Academic researcher
- Civil servant
- Professional services staff
- Other

Approximately how many years experience do you have in your role?

- 0-4 years experience
- 5-9 years experience
- 10+ years experience

If you feel comfortable doing so, please provide an overview of the work you did during the COVID-19 response.

Examples of this might include:

- facilitating meetings
- data management
- producing routine estimates for surveillance and submitting them to advisory bodies.
- developing tools and methods and support their use by other responders
- providing scenario estimates in response to policymakers requests
- writing reports
- academic paper writing
- attending meetings
- supporting researchers
- speaking to the media, … etc.

On a scale from 0 to 10, how would you rate your professional experience in responding to COVID-19 in the UK?

0 - extremely negative

5 - neutral

10 - extremely positive

What professional costs, if any, did you experience from being involved in the response? Think about both short- and long-term costs

What professional benefits, if any, did you experience from being involved in the response? Think about both short- and long-term benefits, for example:

1. number of first author papers
2. promotions/career progression
3. successful grants

On a scale from 0 to 10, how would you rate your personal experience in responding to COVID-19 in the UK?

0 - extremely negative

5 - neutral

10 - extremely positive

What personal costs, if any, did you experience from being involved in the response? Think about both short- and long-term costs

What personal benefits, if any, did you experience from being involved in the response? Think about both short- and long-term benefits

Has your employer or the wider community taken action to help mitigate any of the personal or professional costs you identified?

1. Yes
2. No
3. Somewhat

Can you identify areas where action has been taken and areas where it has not?

Do you think there were barriers to doing effective and sustainable COVID-19 outbreak response work?

1. Yes
2. No
3. Maybe

What were some of the barriers to doing effective and sustainable outbreak response work?

For example, thinking about:

1. funders
2. employer organisations
3. supervisors
4. peers
5. computing resources
6. human resources

What can be done to reduce barriers and better support those involved in outbreak response work in the future?

What were some of the things that helped assist you to do effective research during the outbreak response?

For example, thinking about:

1. funders
2. employer organisations
3. supervisors
4. peers
5. computing resources
6. human resources

Do you think sufficient action is currently being taken to improve future outbreak responses to the standard you think is acceptable?

- Yes
- No
- Unsure

Is there anything else you’d like to add?

### Snapshot questions

Whiteboards with the following questions were available for the first hour of the session for participants to add to with either post it notes or sticker dots. Numbers in brackets [#] represent the number of sticker dots placed by participants.

1. Do you think sufficient action is currently being taken to improve future outbreak responses to the standard you think is acceptable?

- Yes [1]
- [18]
2. Who is responsible for ensuring people are supported, and appropriately credited for their work?

- You [9]
- Your line manager [16]
- Your research team [13]
- Your institution [6]
- Academic funders [5]
- The system more generally [8]
- Other? [0]
3. Summarise your pandemic experience in one word.

- Ambiguous
- Engrossing
- Hard
- Intense (+1)
- Frustration
- Austere
- Valuable (+1)
- Repetitive
- Lonely
- Exhausting
- Focused
- Stressful (x2) (+1) (+1)
- Exciting
- Unprecedented
- Surreal
- Hectic
- Harrowing (+1)

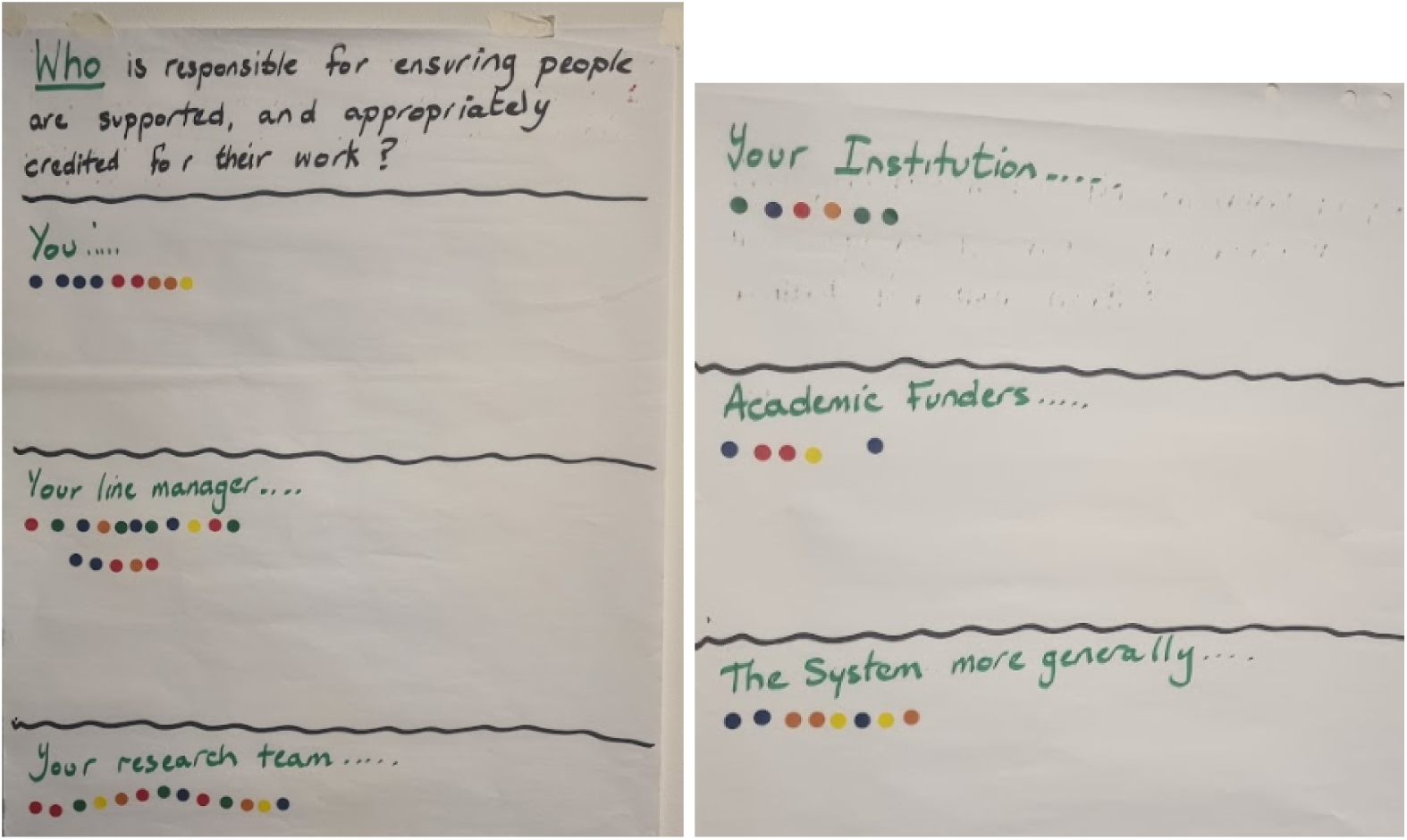

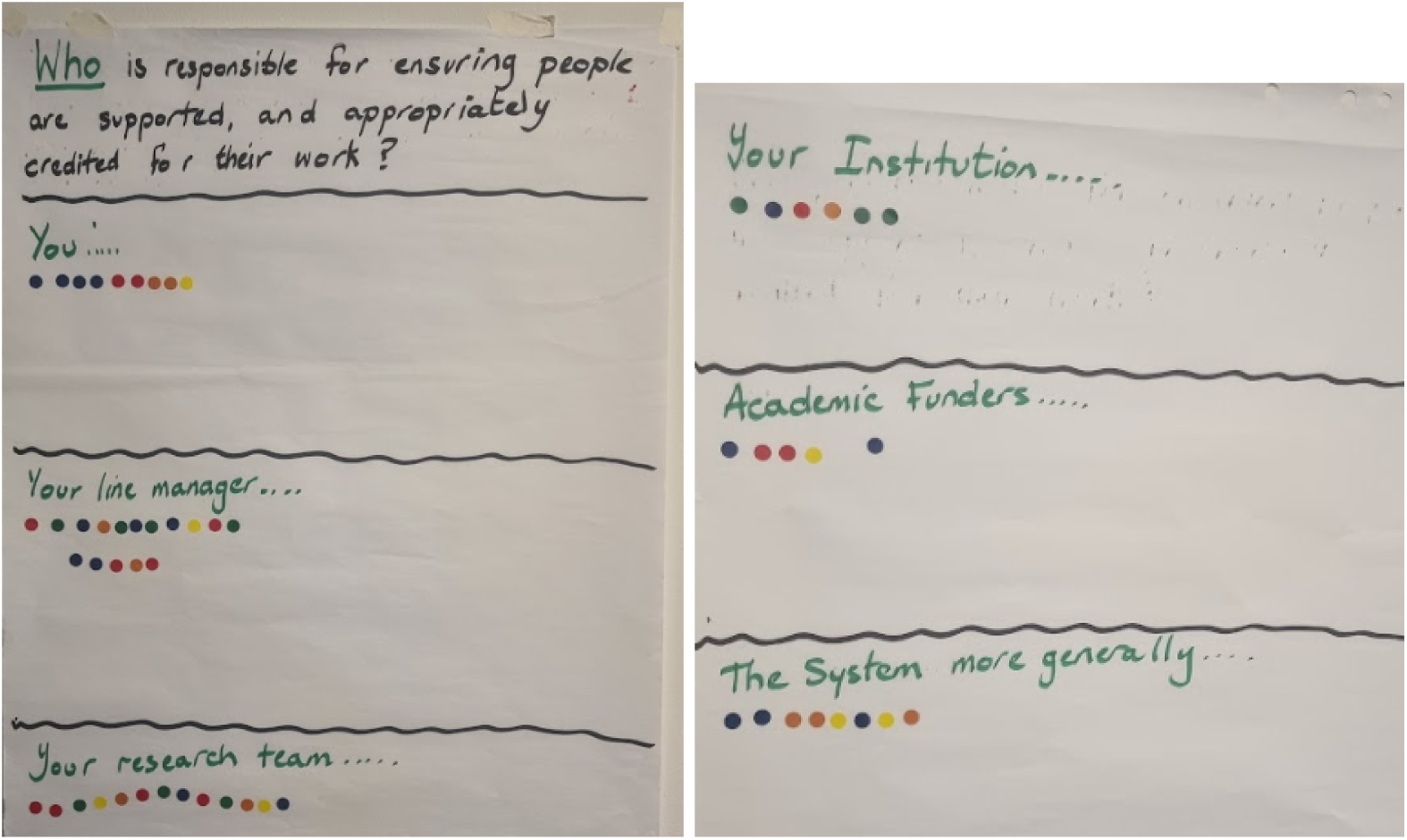

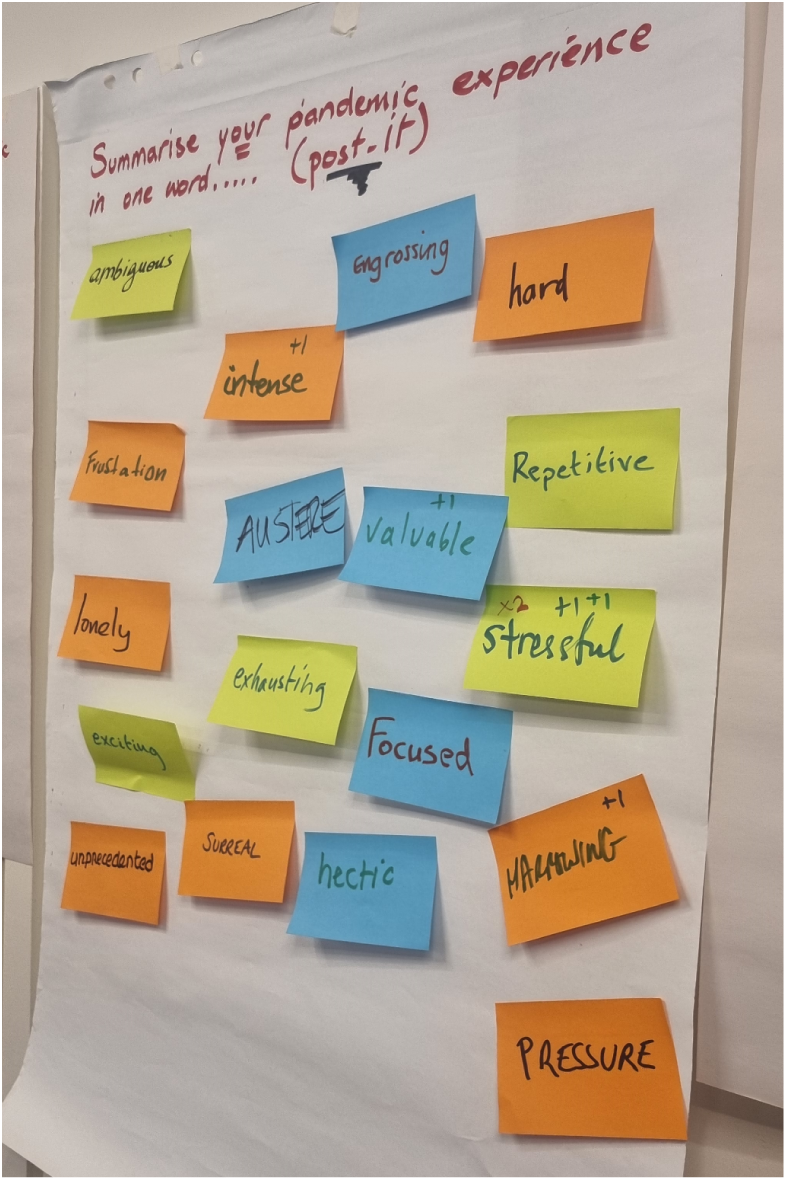

### Session questions

1. What was your pandemic timeline? What were the highs and lows?
2. What was your experience of pandemic work like? Below are some suggested directions to take this:

- What did you expect to experience when you started working on the pandemic response, and how did this differ from what you actually experienced?
- Did you get any benefits?
- Were there any negatives?
- How did your professional and personal experiences differ?
- Do the professional gains outweigh the personal costs or vice versa?
- Do you think privilege played a role in your experience?
3. What were some of the things that helped assist you to do effective research during the outbreak response? Some examples could be:

- Management practices
- Contract duration
- Professional services staff
- Funders
- Employer organisations
- Supervisors
- Peers
- Computing resources
- HR policies

1. Do you think team science was appropriately supported over the pandemic? Some potential directions:

- What elements of team science were most important to your work?
- Has anything changed since prior to the pandemic in terms of support for team science?
- What role did professional services staff, research administrators and managers, press and media teams - have on your ability to do effective response work?
2. Has your employer or the wider community taken action to help mitigate any of the personal or professional costs/challenges you identified? What more can be done?
3. Do you think there were barriers to doing effective and sustainable COVID-19 outbreak response work? If so, what were they?
4. What has been done and what more can be done to reduce any barriers to effective outbreak response work in the future

### Group discussion themes

Participants formed six groups and were asked to summarise their pandemic experience using post it notes, organised into self-identified themes on a whiteboard (one per group). All text is reproduced below, source data photos available on request.

#### Group 1

- Life outside

- Partner in field or not
- Personal circumstances
- Comradeship - shared experience
- Importance of camaraderie in fields like military response - conflicts w/ academic reward structure
- Rewards

- Rewards not attributed equally
- Institutions got awards, not individuals (not all key players)
- Don’t have good mechanisms for rewarding teams (as opposed to individuals)
- Work pressure

- Work life balance
- Overwork
- Lack of strong management
- Lack of leadership
- Line ma!!!1naging
- Mismatch org - task
- Bad stuff

- Authorship issues
- Professional conflicts
- Professional character assassination
- Access & privileges

- Different layers of privilege
- Institutional size (bigger uns = easier)
- Country of origin
- Well connected researcher = better access to data

#### Group 2

- Career direction

- Responsibility at expense of research breadth
- Need to get back to long term planning
- Some career opportunities (but added stress)
- Missing out on non-publication experiences
- Big picture vs detail
- Work process

- Loose in a tornado
- Lots of competing demands
- Difficult to stop rapid work
- Outsourcing prioritisation
- Hybrid work
- (Negativity about) positives

- Shame associated with “doing well”
- Positivenetworking + connections
- Career advancement
- Acknowledge previous bad work habits
- Work “feeling” / support

- Not recognising MH impacts when you’re in it
- Lack of understanding from non C19 colleagues
- Missing “in person” cues in interaction
- Need for more support - burnout
- Lack of support / less guidance
- Missing informal “check-ins” working remotely
- Pride in overwork
- Personal

- Better boundaries / priorities
- Paused personal commitments
- “Is it still March 2020?”

#### Group 3

- A

- Team spirit

- New vs existing
- Lack of support
- Individual shared
- Competitive
- Collaboration
- Institutional

- Leadership
- Protection
- Recognition
- Junior members
- B

- Mental health

- Lack of desire
- Reintegration
- Uncertainty
- Work-life balance
- Children, partners / home, family
- Travel
- Guilt
- Feeling stuck in time-space-role
- Variable speed of time
- Dealing with losses
- Societal impact

- Responsibility
- Value of work
- Representation of work
- Sustainability
- Guilt

#### Group 4

- COVID modellèr specific

- Everyone busy = harder to get support
- Talking about & acknowledge the relative difficulties of each others experiences
- Surreal level of public + media interest (good or bad)
- Differences in levels of success & contribution resulting in inequality in various forms, i.e. lots of success based partly on luck
- Hard to onboard and shift responsibility (sorry new people!)
- Difficult to balance routine modelling and interesting innovative work
- Being lumbered with “shit” jobs
- Worked too much because too much work to do
- More general experiences

- Worked too much because nothing else to do
- Work + the world were one and the same - neither was an escape from the other
- Pressure came in waves “not again…”
- And coincided with Xmas
- Feels like time didn’t exist i.e. no memories for a lot of it
- Really hating the policies for young people

#### Group 5

- Work

- Lack of operational data
- What is “good enough”?
- Not being able to say no
- What was the point of some of the work
- Other work pressures e.g. teaching
- Waves of work
- Working overnight
- Clashes over motivation/priorities
- Emotions

- Powerlessness
- Sadness
- Frustration + anger
- Uncertainty
- Desensitised
- Pride
- Life

- Skewed sense of time
- Life events
- Child care (x2)
- False memories
- What to do next?

#### Group 6

- Structures / work

- Shiqft from egalitarian structure to pre existing hierarchies
- Collaborative work excitement
- Team work challenges
- Gambling / return on investment
- All a blur
- Survivor bias
- Responsibility of managers to look after - who is responsible?
- Trust
- Structures / life

- Burn-out creeps up on you - how to avoid
- Life events (juggling)
- Self-care
- Coping mechanisms
- Personal pandemic preparedness
- Caring for others
- Impact of team members’ personal challenges on everyone
- Emotions / work

- Reconciling different perspectives on hierarchy & team working
- Who gets credit
- Space for communicating
- Guilt
- Comparing self with others
- Frustration
- Feeling useful
- Feeling of lack of legitimacy
- Pride
- Self perception vs others’ perception
- Emotions / life

- Trading off career vs health / everything “just 1 more”
- Personal responsibility (feelings of)
- Maternity leave
- “Fresh blood” (return from leave)
- Fear of missing out

### Themes

Participants were tasked with grouping top-level themes from the group discussions into broad categories as a spider-map. This created the following:

- Work life

- Life outside
- Life
- Personal
- More general experiences
- Work - covid modeller specific

- Work

- Work process
- Work pressure
- Social impact
- Team spirit

- Work “feeling support”
- Mental health
- Emotions
- Rewards

- Structures
- Institutional
- Access & priveleges
- Career direction

- Bad stuff
- (Negativity about) positives

### Recommendations

We reproduce text from the poster boards. We include the implementor suggested by contributors at the end of each statement where this was done.

#### Social impact

- Educate public / policy maker of modelling knowledge for better communication
- Develop primers/training and build on existing links [*research team*]
- Ensure impactful COVID work understood at funder/institution level (esp. If less obvious/visible)
- Support for dissemination of work/case studies [*team/institution/funder*]
- Funder buy-in: creating opportunities to support consolidation of developed methods and tools
- See credit [and fund] all outputs - incl. Software and communication, media, public engagement etc. [*funders/managers/institution*]
- Formalise connections between teams/disciplines/functions - make sure we retain “what worked”
- Provide support (e.g. MH) to reduce burnout + increase retention of institutional memory
- Do[…?]ting connections [*line managers*]

#### Mental health

- Culture shift

- Reframe excellence
- Raising awareness (literacy) - also of team role in crisis response
- Preparedness; peace time team building / leadership training [*wellbeing manager, centre manager*]
- Institutional safeguarding
- “We think we do this well, but we don’t” needs embedding over a sustained time
- Departmental leadership buy in; lead by example (processes - not emailing at night etc) [*funders, institution, people*]
- Resilience training (peers support) [*institution*]
- Psychological first aid *[institution*]
- Individual + management training
- Adequate resourcing of central services + research teams to mitigate against burnout = funding. [*funders, institution*]

#### Career direction

- Now

- Possibly greater acceptance of portfolio PHDs
- Leeway in examiners for PhDs during covid/emergencies
- More recognition of non-traditional outputs
- Future

- Improve ease of movement in/out of public health agencies (eg UKHSA), eg dual positions
- Shifting routine data analysis to public health bodies ASAP
- Stop/end routine activities as soon as they’re not useful
- Find a way to credit confidential work
- How/who

- Institutions & senior academics to write letters of support for PHD students & staff with ‘non-traditional’ outputs [*managers, institution*]
- Joint appointments at public health agencies [*funders*]
- Credit all outputs: academic, tool development, communications, public engagements, confidential reporting
- Clear expectations for promotion [*managers, funders, institutions*]
- Distinguish “academic” vs “emergency” response
- Disaster roster + exercise

#### Work process

- Capacity for cycling between response & research
- Reduce structural reliance on 1-2 people [in] a team performing specific task
- [O]n call system?
- [A]nnual leave
- Prioritise capacity
- Project manager incorporated in team [*department, funder*]
- More transparency from government committees so groups without people on them didn’t get as left behind as it seemed (to me at least)
- Transparent preparedness plan *[UKHSA, govt]*

- Who sits on gov committees
- What their roles/responsibilities
- White paper [*dedicated working group incl funders*]
- Clarify roles for pandemic response: software engineering, policy-related roles
- Broader reward system, e.g. [*funder*]

- Code/software
- reports/briefing notes
- “Middle” authors
- Data collection
- Automation / routine vs. one-off - value/impact?
- Regular re-assessment of cost-effectiveness of tasks
- Ask yourself if you really need to do this so often/ so quickly
- Weekly discussions: priorities [*team leads*]

#### Life/personal

- Mechanism for feeding back & instituting good working practices
- “GWP” [good working practices] reps & a committee that reports to univ./institution executive board

- Identify people to take on the role of rep
- Regular surveys to gauge feelings & elicit suggestions
- Time off in lieu (mandatory)
- Paid overtime (capped)
- Establish working group to make recommendations at a national level [*institution/funders*]
- Realistic funding <> deliverables
- Hobby
- Normalise taking annual leave
- Respect (enforce) working hours [*individuals & institutions*]
- Role models
- Don’t try very hard (Relax)

- Active role
- Guidelines for line managers = cultural change
- Delay emails on weekends and 7pm-7am (allow control)

- Auto-send emails to remind people to take break from work
- PDR [performance and development review] for work/life balance (part of PDR but checked by welfare manager)

- Palm trees on slack
- Normalise people taking holidays > holiday snaps
- “Covid impact statement” but for more general issues
- Have paid wellbeing officer(s)/manager to check in on personal issues & balance, give professional advice, etc…

#### Institutional

- Credit/rewards

- Clear comms on processes
- Meetings non-science (coping, emotion, credits etc)

- > mandatory incorporation in protocols; normalising [*institution*]
- Starter pack

- Processes
- Support & social
- Technical
- Mentorship/buddies
- Available training
- survey input regular
- share experience between institutions [*department groups*]
- Resilience

- Firefighter mentality
- Back-up / replaceable

- learn from other professions
- Structure ahead of time & practice [*dedicated working group*]
- Responsibility

- Hierarchy
- Map roles
- Back-up/replaceable
- Function of/process for support roles (software, admin/comms)

- Starter pack
- available guidelines
- structured updating [*institution, department*]
- Clear guidance

- Communication
- Expectations
- Reference resource

- Project manager
- career path for support roles with growth
- comms / software / admin project management [*funder, institution*]
- Manager training

- Niche to normal
- Timely, frequent
- across seniority

- Mandatory training [*institution*]
- Lessons learned

- Exit interview feed into updating processes
- Left since start of pandemic
- All staff, not only academic

- Survey / exit interview [*manager/department*]

#### Additional

- Pay rise
- Now now v just now prioritisation

**Figure 1.**
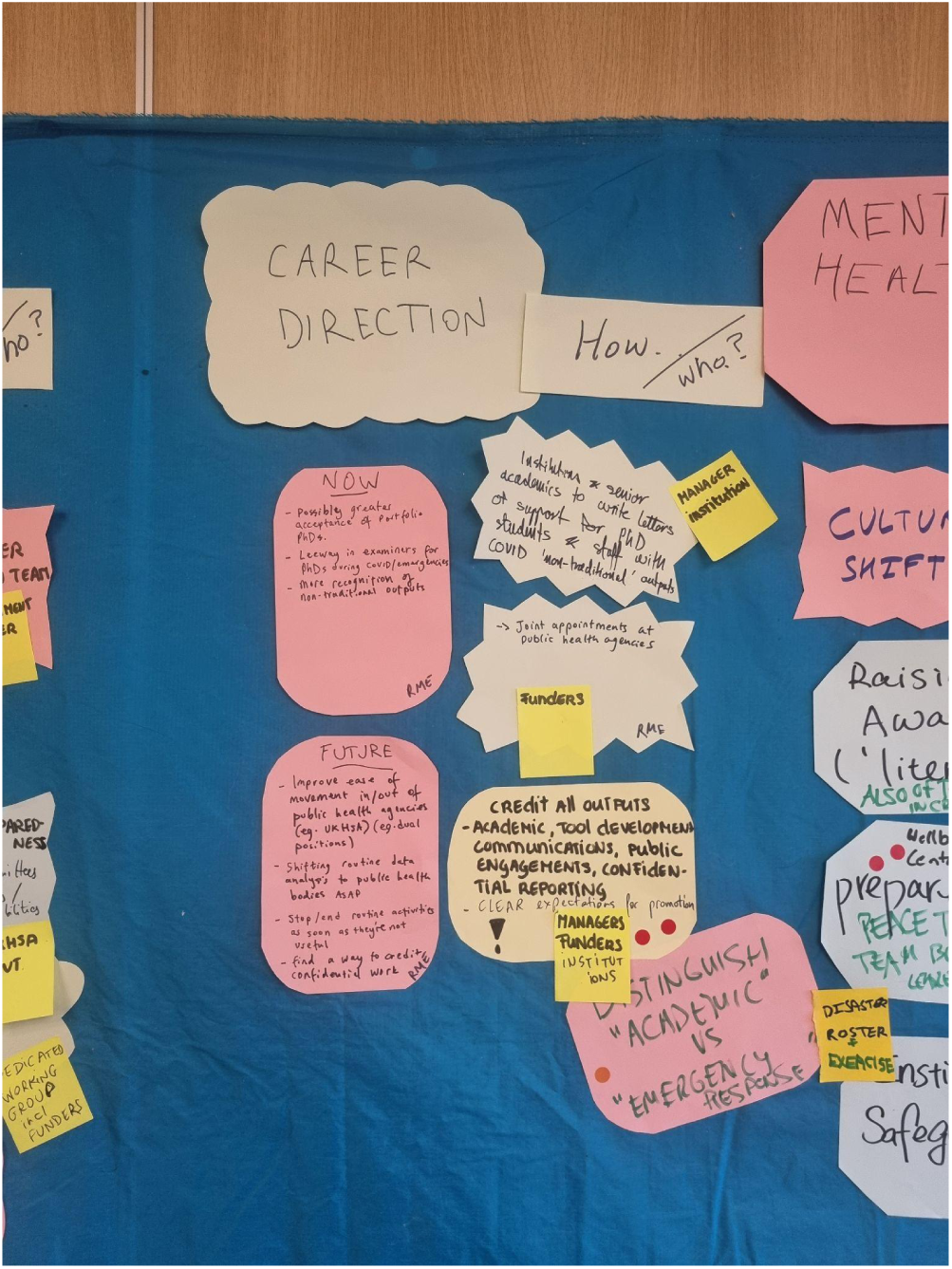
Example of board showing recommendations

### Priority recommendations

These recommendations were highlighted by at least one participant during a group discussion. Recommendations which were highlighted by two or more participants are shown in bold.

1. Research teams should ensure that impactful COVID work is understood by funders and institutions, especially if less obvious/visible.
2. Research teams should initiate sustainable team building and training programs during non-response periods.
3. Employers should ensure sustainable funding for academic and professional services roles to reduce burnout risks.
4. Employers should develop processes and guidelines for career growth support for professional services staff.
5. Employers need to create or expand Wellbeing Officer roles to monitor work-life balance and provide guidance.
6. Incorporation of work-life balance components in annual performance and development reviews is essential for employers.
7. Employers should implement capped paid overtime and formal Time Off In Lieu policies and routinely analyse and act upon staff survey and exit interview data.
8. Funding bodies should adjust eligibility criteria to adequately compensate non-academic staff for activities.
9. Funding bodies should refine impact measures to acknowledge all outputs, including academic, tool development, communications, public engagement, and confidential reporting.

